# Admixture with cultivated sunflower likely facilitated establishment and spread of wild sunflower (*H. annuus*) in Argentina

**DOI:** 10.1101/2024.02.22.581669

**Authors:** Fernando Hernández, Román B. Vercellino, Marco Todesco, Natalia Bercovich, Daniel Alvarez, Johanne Brunet, Alejandro Presotto, Loren H. Rieseberg

**Affiliations:** Department of Botany and Biodiversity Research Centre, University of British Columbia, Vancouver, British Columbia, Canada; Departamento de Agronomía, CERZOS, Universidad Nacional del Sur (UNS)-CONICET, Bahía Blanca, Argentina; Michael Smith Laboratories, University of British Columbia, Vancouver, Canada; Irving K. Barber Faculty of Science, University of British Columbia Okanagan, Kelowna, Canada; Estación Experimental Agropecuaria INTA Manfredi, Manfredi, Córdoba, Argentina; Vegetable Crops Research Unit, USDA-ARS, Madison, WI, USA

**Author notes:** **Correspondence** Fernando Hernández, Department of Botany and Biodiversity Research Centre, University of British Columbia, Vancouver, British Columbia, Canada.

## Abstract

A better understanding of the genetic and ecological factors underlying successful invasions is critical to mitigate the negative impacts of invasive species. Here, we study the invasion history of *Helianthus annuus* populations from Argentina, with particular emphasis on the role of post-introduction admixture with cultivated sunflower (also *H. annuus*) and climate adaptation driven by large haploblocks. We conducted genotyping-by-sequencing of samples of wild populations as well as Argentinian cultivars and compared them with wild (including related annual *Helianthus* species) and cultivated samples from the native range. We also characterized samples for 11 known haploblocks associated with environmental variation in native populations to test whether haploblocks contributed to invasion success. Population genomics analyses supported two independent geographic sources for Argentinian populations, the central United States and Texas, but no significant contribution of related annual *Helianthus* species. We found pervasive admixture with cultivated sunflower, likely as result of post-introduction hybridization. Genomic scans between invasive populations and their native sources identified multiple genomic regions with evidence of selection in the invaded range. These regions significantly overlapped between the two native-invasive comparisons and showed disproportionally high crop ancestry, suggesting that crop alleles contributed to invasion success. We did not find evidence of climate adaptation mediated by haploblocks, yet outliers of genome scans were enriched in haploblock regions and, for at least two haploblocks, the cultivar haplotype was favored in the invaded range. Our results show that admixture with cultivated sunflower played a major role in the establishment and spread of *H. annuus* populations in Argentina.

## INTRODUCTION

Invasive species represent a major threat to local biodiversity (Mollot et al., 2017), and to crop and pasture production when they invade agricultural habitats (Kuester et al., 2014). Invasives can hybridize with their cultivated relatives, resulting in the introduction of herbicide and pesticide resistance into potential agricultural weeds and limiting the efficacy of herbicide and pest control technologies (Ellstrand et al., 2010; Vercellino et al., 2023a). Invasive species are also excellent model systems to study rapid adaptation, as in many cases they were introduced to novel environments and successfully established and spread within decades or a few centuries (Prentis et al., 2008; Bock et al., 2015; Colautti et al., 2017).

Invasive species are often non-natives and a central goal of invasion genomics is to identify the genetic and ecological factors underlying the successful establishment and range expansion of introduced species. However, the genetic make-up of invasive species can be strongly influenced by stochastic processes such as the non-random introduction of genetic variation from the native range, founder effects during introduction, allele surfing during range expansion, and, in the cases of multiple introductions, admixture between historically isolated populations (Dlugosch and Parker, 2008; Dlugosch et al., 2015; Schrieber and Lachmuth, 2017). These processes complicate the identification of loci underlying invasion success and, for this reason, an understanding of the invasion history should guide evolutionary inferences.

During introduction, non-native species experience strong demographic bottlenecks, which are expected to reduce their genetic diversity, fitness, and evolutionary potential (Dlugosch and Parker, 2008; Keller and Taylor, 2008; Sherpa and Després, 2021). Yet some of these species successfully establish, persist, and become invasive, a dilemma referred to as the genetic paradox of invasions (Estoup et al., 2016). Phenotypic plasticity, defined as the ability of a genotype to express different phenotypes in different environments (Baker, 1965; Richards et al., 2006; Liao et al., 2016), and prior adaptation, where locally adapted populations are introduced to regions with similar environmental characteristics (Liu et al., 2004; Hufbauer et al., 2012), can explain this genetic paradox, as neither of these mechanisms involve evolutionary changes in the invaders. However, introduced populations often show similar (or even higher) genetic diversity relative to their native counterparts (Dlugosch and Parker, 2008; Dlugosch et al., 2015; Estoup et al., 2016), and rapid evolution has been hypothesized as a major factor contributing to successful invasions (Felker-Quinn et al., 2013; Turner et al., 2013; Oduor et al., 2016; Barker et al., 2017; O’Neill et al., 2017).

Several studies have shown an association between admixture and invasion success (Hovick and Whitney, 2014; Keller et al., 2014; van Kleunen et al., 2015; Vilatersana et al., 2016; Barker et al., 2017; van Boheemen et al., 2017). Admixture may facilitate establishment and spread by alleviating demographic and genetic bottlenecks during the earliest stages of invasion, augmenting the genetic diversity of introduced populations, increasing the fitness of hybrids due to heterosis, and creating novel genetic combinations on which selection can act (Rieseberg et al., 2003; Dlugosch and Parker, 2008; Dlugosch et al., 2015). Some introduced wild species can also hybridize with their cultivated relatives, leading, in some cases, to the origin of aggressive agricultural weeds (Campbell et al., 2009; Ellstrand et al., 2010; Pandolfo et al., 2016; Le Corre et al., 2020; Vercellino et al., 2023a).

Rapid evolution in the introduced range is expected to occur through standing genetic variation rather than new mutations, and large effect loci are expected to play a major role (Battlay et al., 2023). Adaptive allele combinations may be maintained and spread in genomic regions of reduced recombination, forming large adaptive haplotypes (hereafter named haploblocks), which are mostly associated with chromosomal inversions (Kirkpatrick and Barton, 2006; Ortiz-Barrientos et al., 2016; Wellenreuther and Bernatchez, 2018; Faria et al., 2019). Haploblocks are associated with environmental adaptation and ecotype formation across the tree of life (Lowry and Willis, 2010; Jones et al., 2012; Jay et al., 2018; Hager et al., 2022), and a recent study showed how such variants have facilitated the adaptation of an invasive species over contemporary timescales (Battlay et al., 2023).

Wild sunflower (*Helianthus annuus*) is native to North America and has been introduced and declared invasive in Australia (Seiler et al., 2007), southern Europe (Muller et al., 2011), Israel (Hübner et al., 2022), and Argentina (Poverene et al., 2008). In Argentina, previous studies based on phenotypic data support a likely introduction of most wild sunflower populations from the central United States (Cantamutto et al., 2010a; Hernández et al., 2019b) and a subsequent spread over the central regions of Argentina via agricultural machinery (Cantamutto et al., 2010b). Argentinian populations resemble populations from their putative geographic source for most phenotypic traits, but introduced and native populations show divergence in ecologically relevant traits like disk size, seed size, and seed dormancy (Hernández et al., 2018, 2019a). Because these ecologically relevant traits have shifted towards the crop phenotypes in the introduced wild populations, and because crop and wild naturally hybridize in Argentina (Ureta et al., 2008), the introgression of crop alleles into wild populations is thought to have facilitated their evolution (Cantamutto et al., 2010a; Hernández et al., 2019b) . However, the permanent introgression of crop alleles in introduced wild sunflower populations, or evidence that such alleles are favored by natural selection, has yet to be demonstrated.

Beside admixture with cultivated sunflower, hybridization with other wild *Helianthus* species must also be considered. Several wild annual species of *Helianthus* (e.g., *H. annuus*, *H. argophyllus*, and *H. petiolaris*) were deliberately introduced in the 1950’s in Argentina for crop breeding purposes (Bertero de Romano and Vázquez, 2003). At least two of these species (*H. annuus* and *H. petiolaris*) have naturalized (Poverene et al., 2008), grow in sympatry in some areas (Cantamutto et al., 2008; Mondon et al., 2018), and may grow in close proximity to cultivated sunflower (Ureta et al., 2008; Mondon et al., 2018). Therefore, it is important to determine not only whether cultivated sunflower, but also other *Helianthus* species (e.g., *H. petiolaris* and *H. argophyllus*), have contributed to the ancestry of Argentinian *H. annuus* populations.

In the native range, 11 large haploblocks are associated with multiple adaptive traits and climate variables, and these haploblocks explain most of the genomic and phenotypic variation detected between the texanus (adapted to warmer and less continental climates) and non-texanus ecotypes (Todesco et al., 2020; Owens et al., 2021). As these haploblocks segregate in the putative geographic source of invasive populations (the central US), we expect that both texanus and non-texanus haplotypes were introduced to Argentina. We further hypothesize that the haploblocks associated with the texanus ecotype may have facilitated the rapid adaptation of wild sunflower populations in central Argentina, which is similar to the climate of the texanus ecotype in its native range (Hernández et al., 2019a).

The specific aims of this study are to 1) identify the native source populations of Argentinian populations; 2) evaluate whether crop introgression influenced post-introduction divergence; and 3) evaluate whether the haploblocks associated with climate adaptation in the native range facilitated the establishment and spread of sunflowers in Argentina.

## MATERIALS AND METHODS

### Plant material

For this study, we extracted DNA from 153 Argentinian samples: 83 individuals from 10 wild *H. annuus* populations collected in disturbed, nonagricultural habitats (mostly roadsides, hereafter refered as ANN INV), 9 individuals from one weedy *H. annuus* population collected within an agricultural field (FERAL), 13 inbred line parents of modern cultivars (CROP MODERN), and 48 samples from old cultivars (open pollinated varieties developed in Argentina and released from 1930 – 1980; CROP OLD) (Table 1). Seeds from the wild and the weedy populations were collected between 2017 and 2022 (Table 1) from at least 50 plants per population, while seeds from modern and old cultivars were generously provided by breeders at ACA Semillas (Argentinian seed company) and INTA Manfredi (Argentinian public institution of agricultural technology), respectively. Seeds were germinated and grown to the seedling stage (up to two leaves) and cotyledons were cut and stored at −20°C until DNA extraction. Genomic DNA was extracted from cotyledons, following a modified CTAB procedure. DNA quantification was performed with a Qubit 2.0 fluorometer and DNA integrity was analyzed by electrophoresis using a 1% agarose gel. The 153 DNA samples were genotyped by RAD-seq following the approach described by (Elshire et al., 2011). Briefly, DNA samples were digested with the restriction enzyme ApeKI, barcoded, and pooled for sequencing using 150 bp paired end reads on an Illumina NovaSeq 6000 platform (500 million reads in total, ∼3 million reads per sample). DNA extractions were performed at the GENeTyC in Bahia Blanca, Buenos Aires, Argentina while library preparation and sequencing were performed at the University of Wisconsin-Madison Biotechnology Center DNA Sequencing Facility. Raw reads of Argentinian samples will be submitted to the NCBI Short Read Archive (SRA) upon manuscript acceptance.

**Table 1.** Information of Argentinian samples sequenced for this study. Sample size (N) after filtering. Lat, Long: latitude and longitude of collection site. BIO1: mean annual temperature (°C), BIO12: annual precipitation (mm).

| Group | Pop ID | Province | N | Lat | Long | BIO1 | BIO12 |
| --- | --- | --- | --- | --- | --- | --- | --- |
| ANN INV | BAR | La Pampa | 7 | -36.17 | -63.87 | 15.93 | 723 |
|  | CAR | Buenos Aires | 7 | -32.27 | -62.92 | 16.84 | 801 |
|  | DIA | Entre Rios | 6 | -32.05 | -60.63 | 18.23 | 1033 |
|  | EBA | Córdoba | 9 | -30.35 | -63.83 | 17.77 | 770 |
|  | JCE | Córdoba | 10 | -33.67 | -63.47 | 16.68 | 876 |
|  | LMA | Mendoza | 9 | -34.93 | -68.23 | 15.04 | 296 |
|  | MAG | San Juan | 10 | -31.95 | -68.45 | 17.89 | 255 |
|  | RCU | Córdoba | 9 | -33.15 | -64.33 | 16.53 | 839 |
|  | VME | San Luis | 10 | -33.87 | -65.05 | 16.64 | 634 |
|  | WIN | La Pampa | 6 | -36.30 | -65.58 | 15.81 | 593 |
| FERAL | BRW | Buenos Aires | 9 | -38.27 | -60.12 | 14.00 | 769 |
| CROP ARG | CROP OLD | - | 48 | - | - | - | - |
|  | CROP MODERN | - | 13 | - | - | - | - |

To identify potential contributions of other annual *Helianthus* species and of cultivated sunflower to the ancestry of the argentinian *H. annuus* populations, we merged the dataset generated for this study with previously generated whole-genome sequence datasets of cultivated (Hübner et al., 2019) and wild native sunflower (Todesco et al., 2020).

### Sample alignment, variant calling, and filtering

Raw reads were demultiplexed with Sabre and trimmed using *Cutadapt* (Martin, 2011) with default options, and mapped to the reference assembly HA412-HOv2 (Huang et al., 2023) using *BWA-MEM* (Li and Durbin, 2009). Variant calling was performed with *samtools* (Li et al., 2009). Then, we used *vcftools* (merge function) to merge our dataset with two previously published whole genome sequence datasets (WGS), comprising 1505 samples from natural populations of *Helianthus* species [299 of *H. argophyllus* (PHY), 401 of *H. petiolaris* subsp. *petiolaris* and *H. petiolaris* subsp. *fallax*, 86 of *H. niveus* (PET), and 719 of *H. annuus* (ANN NAT and ANN TX)] (Todesco et al., 2020), and 287 cultivars comprising the sunflower association mapping (CROP SAM) population, which represents all major genetic groups within the cultivated germplasm (Hübner et al., 2019). Reads from all samples were mapped to the same reference assembly (HA412-HOv2). Information on the samples included in each dataset can be found in Table S1. To avoid biases due to the broader coverage expected in the WGS dataset compared to the RAD-seq one, the variant call format (VCF) file with WGS samples was filtered to retain only sites present in at least 70% of the RAD-seq samples before merging. The resulting VCF files were filtered to retain only biallelic SNPs with MAF> 0.01, minimum quality> 30, and present in at least 70% of the samples. The final VCF file will be deposited in the DRYAD repository upon manuscript acceptance. Two datasets were produced, dataset 1 contained all *Helianthus* species: 1947 samples and 6,094 SNPs, and dataset 2 with only *H. annuus*: 872 samples and 24,919 SNPs (Table S1).

### Climatic characterization of native and invaded ranges

Nineteen biologically relevant climatic variables were obtained from the WorldClim2 dataset (Fick and Hijmans, 2017) for each of the 81 wild *H. annuus* populations (Table S2). We ran pairwise correlation analyses for all the variables and, for all the pairwise comparisons with r> [0.8], one variable of each pair was removed from the analysis to avoid redundancy. After this removal, 12 BIOCLIM variables represented the climate input data for both principal component and clustering analyses.

### Population genomics and admixture

For population structure analyses (PCA, DAPC, and ADMIXTURE), the two datasets were filtered to retain only SNPs in low linkage disequilibrium (LD; r^2^< 0.2) with the R package *SNPRelate* (Zheng et al., 2012). PCA and DAPC were performed with the R packages *SNPRelate* and *adegenet* (Jombart et al., 2010), respectively, and ADMIXTURE analysis with *admixture v.1.3* (Alexander and Lange, 2011) with K=2 - 10 and cross-validation (cv)= 10. Pairwise F_ST_ between populations and between genetic groups identified with DAPC were estimated with *Arlequin v.3.5.2.2* (Excoffier et al., 2005).

To identify the geographic origin of invasive populations, we used a deep learning method implemented in *Locator* (Battey et al., 2020). The 719 samples of wild native *H. annuus* (including both texanus and non-texanus ecotypes) from 71 geo-referenced native populations were used to predict the origin of the 83 wild *H. annuus* Argentinian samples using the low LD dataset. We excluded the nine samples from the weedy population (BRW) because of its known recent feral origin in Argentina (Hernández et al., 2022). Argentinian samples were not assigned to populations in this analysis.

### Crop introgression

In order to maximize the number of SNPs, crop introgression analyses were performed with dataset 2, which only includes *H. annuus* samples. First, we used admixture results at K= 2 to measure the proportion of the cultivated cluster in native and invasive samples and estimated the average proportion for each population. To test for crop to wild introgression post introduction, we performed ABBA-BABA tests across four populations: P1, P2, P3, and the outgroup (O), related by the phylogeny (((P1, P2), P3), O). The allele carried by the outgroup is designated as the ancestral allele (A), while the derived allele is designated as B. In all tests, individuals from native populations were used as P1, individuals from invasive populations were used as P2, all cultivated samples from Argentina were used as P3 (n= 61), and one sample for each of four perennial sunflower species (*H. decapetalus, H. divaricatus, H. giganteus,* and *H. grosseserratus*) were used as the outgroup. Under the null hypothesis, which assumes no gene flow after the P1 (native) and P2 (invasive) split, the ABBA (B shared by P2 and P3) and BABA (B shared by P1 and P3) patterns are expected to occur with equal frequency due to incomplete lineage sorting. A significant increase in ABBA or BABA is consistent with introgression between P3 and either P1 (ABBA < BABA, negative D values) or P2 (ABBA > BABA, positive D values). We performed one test for each of the 10 argentinian populations and one test with all argentinian samples as P2. Trios showing Z-scores > 3 were considered statistically significant. ABBA-BABA tests were performed using *Dsuite* (Malinsky et al., 2021).

### Haploblock genotyping

Coordinates of structural variants reported in Todesco et al., (2020) were used to assign SNPs to 11 known haploblocks. For each haploblock, a supervised machine learning approach was used to determine the genotype of 258 samples (including all wild and cultivated Argentinian samples; test dataset) using 614 native samples with known haploblock genotypes as a training dataset (Table S3). The haploblock genotypes of the training samples were used as the categorical response variable (three genotypes: homozygotes for either haplotype or heterozygotes) while the SNPs within each haploblock (which varied from 39 to 692) were used as predictor variables. Random forest models were constructed using the R package *randomForest* (Liaw & Wiener, 2002). The number of decision trees was set to 500, and confusion matrixes were used to estimate the classification error (percent of misclassified genotypes). Trained models were then used to infer the haploblock genotypes of the samples included in test dataset.

### Genome scan and outlier detection

We performed a genome scan as implemented in *Bayescan v.2.1* (Foll and Gaggiotti, 2008) to identify outliers with high F_ST_ between two *a priori* defined groups (native source and invasive populations). SNPs with MAF< 5% and missing data> 30% were filtered out. The analysis was run with default settings, which included 20 pilot runs of 5000 steps each, followed by 50000 burn-in and 5000 sampling steps with a thinning interval of 10, and the prior odds parameters set to 10. SNPs with q-value< 5% for the model including selection were retained as outliers.

## RESULTS

### Population structure

With all *Helianthus* species (dataset 1), PCA and DAPC suggested four main genetic groups (Fig. 1A and 1B), largely corresponding (> 99%) to samples of 1) *H. petiolaris* subsp. *petiolaris*, *H. petiolaris* subsp. *fallax*, and *H. niveus* subsp. *canescens* (PET); 2) *H. argophyllus* (PHY); 3) wild *H. annuus* (ANN TX and ANN NAT for texanus and non-texanus ecotypes, respectively); and 4) cultivated *H. annuus* (CROP SAM). All invasive samples (ANN INV) but one (which was assigned to the cultivated group), were assigned to the wild *H. annuus* genetic group while all cultivated samples from Argentina, either old or modern (CROP ARG) and samples of the feral population (BRW) were assigned to the cultivated *H. annuus* genetic group (Table S1).

**Figure 1.**
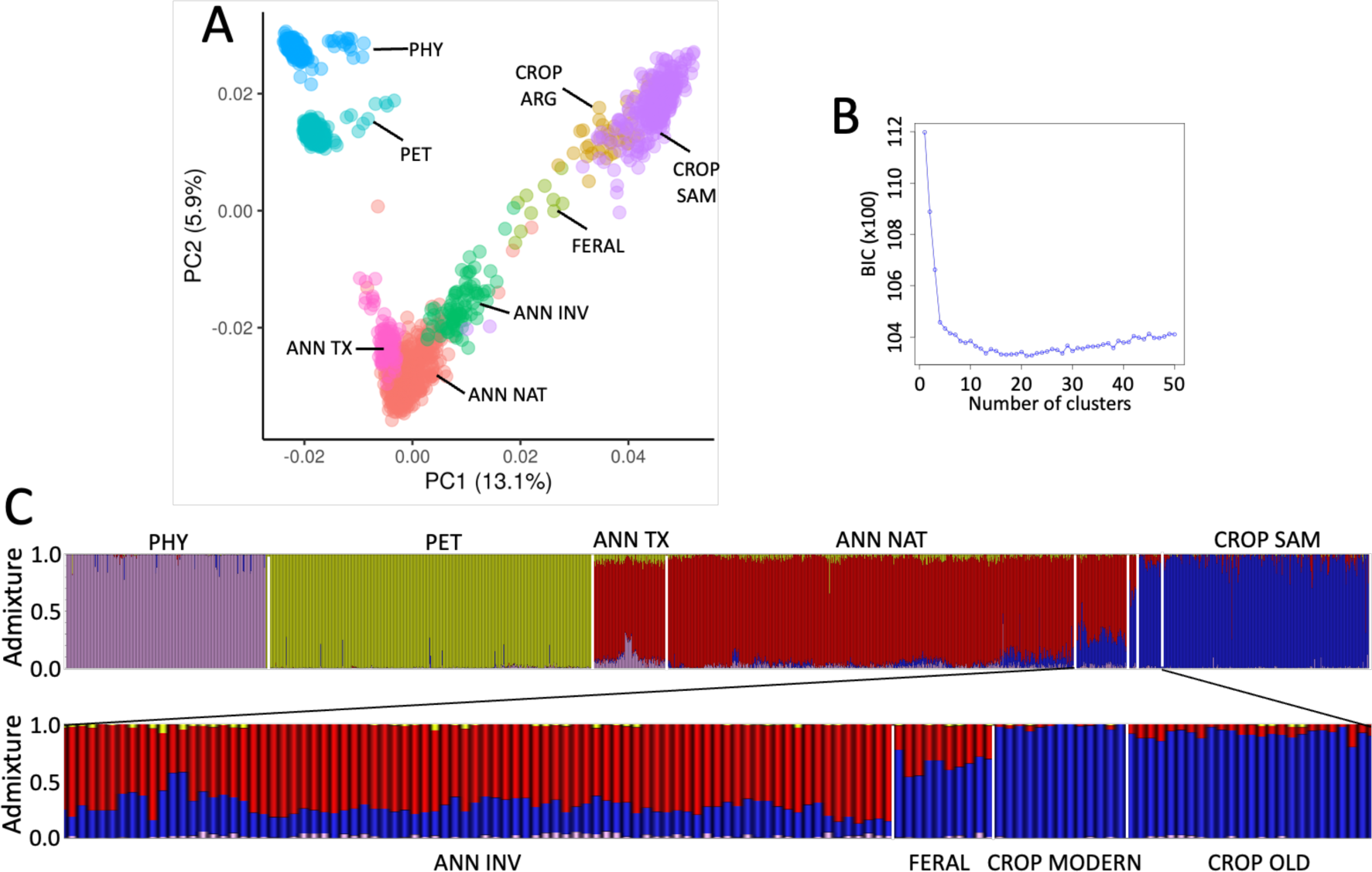
Population structure. A: Principal component analysis (PCA). B: Discriminant analysis of principal components (DAPC). C: ADMIXTURE at K= 4. PHY: *Helianthus argophyllus*; PET: *H. petiolaris* and *H. niveus*; ANN TX: native *H. annuus*, texanus ecotype; ANN NAT: native *H. annuus*, non-texanus ecotype; ANN INV: Argentinian invasive *H. annuus*; FERAL: feral population from Argentina (BRW); CROP ARG: modern and old cultivars from Argentina (CROP MODERN and CROP OLD); CROP SAM: cultivars from the sunflower association mapping population.

Using ADMIXTURE, at K= 4, samples split into the same four groups, and invasive samples showed large proportions of crop ancestry (Fig. 1C), ranging from 0.11 to 0.56 (mean= 0.26). Although *H. petiolaris* and *H. argophyllus* contributed to the ancestry of some native populations (e.g., PHY introgression into ANN TX), their contribution to the ancestry of invasives was negligible (mean= 0.01 and 0.019 for PET and PHY ancestries, respectively) (Fig. 1C). Therefore, to maximize the number of SNPs in our dataset, further analyses were performed on all native (ANN NAT= 609 and ANN TX= 110), invasive (ANN INV= 83), and feral (FERAL= 9) *H. annuus* samples, and on cultivars from Argentina (CROP ARG= 61) (dataset 2) (872 samples and 24,919 SNPs).

### Geographic source of invasive populations

An LD-pruned set of 719 samples from 71 geo-referenced native *H. annuus* populations (ANN NAT and ANN TX) was used to predict the origin of the 83 wild *H. annuus* Argentinian samples (ANN INV). *Locator* predicted that most Argentinian samples (67/83 samples, 8/10 populations, hereafter INV1, n= 67) were introduced from a region in central United States, ranging from latitude 38.3 to 44.7 and longitude -99.3 to -96.0 and comprising parts of the states of Kansas, Iowa, Nebraska, and South Dakota (Fig. 2A and 2B). In addition, samples of two other Argentinian populations (DIA and MAG; hereafter INV2, n= 16) were predicted to have been introduced from Texas (Fig. 2A and 2C). Therefore, we assumed two independent geographic sources for wild Argentinian populations: the first one in the central US (NAT, n= 69), represented by six populations (Barton, Norton, and Phillips from Kansas, Mills and Woodbury from Iowa, and Union from South Dakota; Fig. 2B) and the second one in Texas (TX, n= 20), represented by two populations (Nueces and Tarrant from Texas; Fig. 2C). Genetic differentiation was significantly higher for comparisons within Argentina than between ranges (Fig. S1), suggesting restricted gene flow in the invaded area.

**Figure 2.**
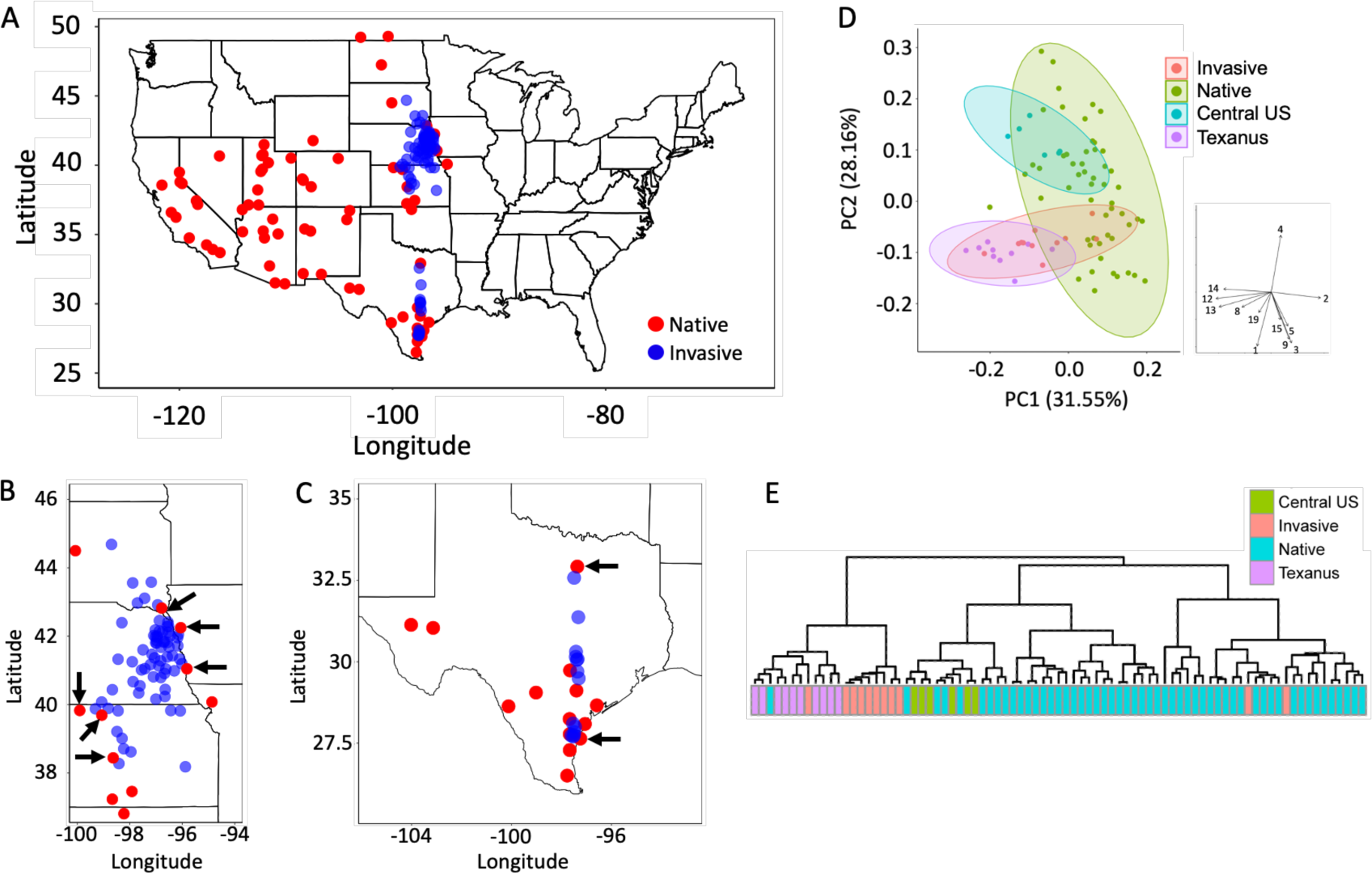
Geographic source of wild Argentinian samples based on Locator and climate characterization of native and invaded ranges. A. Map showing all native populations (red dots, each represented by 10 samples) used in Locator to predict the geographic source of invasive samples (blue dots). B and C: Central US and Texas regions where most Argentinian samples are predicted to come from. Arrows indicate populations used as native sources hereafter (NAT in B and TX in C). D and E: PCA and clustering analyses based on 12 BIOCLIM variables with low correlation (r< [0.8]). In D and E Native refers to native populations other than Texanus and Central US populations.

In the PCA with the 12 climate variables, the first two components captured 60% of total variation (Fig. 2D). PC_1_ was negatively associated to precipitation while PC_2_ was negatively associated to temperature and positively associated to continentality (Fig. 2D). The distribution of invasive populations largely overlapped with the Texanus ecotype (Fig. 2D). Similarly, in the clustering analysis, all invasive populations but two clustered with Texanus populations, and none of them clustered with populations from the putative geographic source (Fig. 2E).

### Crop introgression

To quantify admixture with cultivated sunflower, we used dataset 2, which included only *H. annuus* samples. In ADMIXTURE, at K= 2, cultivated and wild native samples were well-separated (mean crop ancestry = 0.998 and 0.039, respectively; Fig. S2), and invasive samples showed large proportions of crop ancestry, ranging between populations from 0.17 in WIN to 0.43 in CAR (mean= 0.30; Fig. S2). Populations from the putative geographic source in the central US also showed significant proportions of crop ancestry, ranging from 0.07 in Barton, Kansas to 0.18 in Mills, Iowa (mean= 0.12), while populations from the putative geographic source in Texas showed low proportions of crop ancestry: 0.01 in Nueces and 0.04 in Tarrant (Fig. S2). When we estimated crop ancestry proportions using dataset 1, which included the CROP SAM samples as well as the other *Helianthus* species, results showed strong correlation with the estimations using dataset 2 (ρ = 0.98 for pooled native source and invasive samples and ρ = 0.96 for invasive and native sources grouped separately).

For ABBA-BABA tests, individuals from native populations from the central US geographic source (NAT) or the Texas geographic source (TX) were used as P1, all samples for each invasive population were used as P2, all cultivated samples from Argentina were used as P3 (CROP ARG, n= 61), and one sample for each of four perennial sunflowers species (*H. decapetalus, H. divaricatus, H. giganteus,* and *H. grosseserratus*) were used as the outgroup. ABBA-BABA tests were statistically significant (Z> 3) for all but one invasive population (WIN; Table 2) and the estimated fraction of introgression (f_4_-ratio) varied from 0.06 in WIN to 0.36 in CAR (mean= 0.25) (Table 2). Pooling samples of the INV1 group and using them as P2 gave similar results, as did the use of pooled samples of the INV2 group as P2 (Table 2).

**Table 2.** Patterson’s D statistic, Z-scores, p-values, and f4-ratios for each invasive population. NAT: 69 individuals from six native populations from the central US (Barton, Phillips, and Norton from Kansas, Mill and Woodbury from Iowa, and Union from South Dakota); CROP: all cultivars from Argentina (n= 61); TX: 20 individuals from two populations from Texas (Tarrant and Nueces); INV1: 67 wild Argentinian individuals from BAR, CAR, EBA, JCE, LMA, RCU, VME, and WIN populations; INV2: 16 wild Argentinian individuals from DIA and MAG populations. One sample for each of four perennial sunflowers species (*H. decapetalus*, *H. divaricatus*, *H. giganteus, and H. grosseserratus*) were used as outgroup. Crop ancestry proportion from ADMIXTURE at K= 2 is included for comparison. Details on the invasive populations are presented in Table 1.

| P1 | P2 | P3 | Patterson's D | Z-score | p-value | $f_4$-ratio | ADMIXTURE |
| --- | --- | --- | --- | --- | --- | --- | --- |
| NAT | BAR | CROP | 0.186 | 11.2 | 2.3E-16 | 0.21 | 0.25 |
| NAT | CAR | CROP | 0.308 | 11.0 | 2.3E-16 | 0.36 | 0.43 |
| NAT | EBA | CROP | 0.167 | 6.1 | 1.1E-09 | 0.19 | 0.25 |
| NAT | JCE | CROP | 0.185 | 11.4 | 2.3E-16 | 0.21 | 0.27 |
| NAT | LMA | CROP | 0.274 | 14.2 | 2.3E-16 | 0.32 | 0.38 |
| NAT | RCU | CROP | 0.186 | 11.5 | 2.3E-16 | 0.21 | 0.26 |
| NAT | VME | CROP | 0.231 | 13.9 | 2.3E-16 | 0.27 | 0.32 |
| NAT | WIN | CROP | 0.055 | 2.9 | 4.1E-03 | 0.06 | 0.17 |
| TX | DIA | CROP | 0.369 | 13.7 | 2.3E-16 | 0.38 | 0.34 |
| TX | MAG | CROP | 0.321 | 12.6 | 2.3E-16 | 0.33 | 0.31 |
| TX | INV2 | CROP | 0.336 | 14.4 | 2.3E-16 | 0.34 | 0.32 |
| NAT | INV1 | CROP | 0.204 | 20.5 | 2.3E-16 | 0.23 | 0.30 |

### Haploblocks genotyping

In our filtered dataset 2 for *H. annuus* (872 samples and 24,919 SNPs), we found SNPs within each of the 11 haploblocks, with the number of SNPs per haploblock ranging from 39 (for haploblock ann14.02) to 692 (for haploblock ann13.01). Random forest models predicted haploblock genotypes with high accuracy: error rates varied from 0% for ann14.01 to 6.8% for ann17.01 (mean= 1.36%). Most of the misclassified genotypes were heterozygotes misclassified as homozygotes or vice versa, while only three individuals (out of 614 per haploblock used in the training datset) that were homozygotes for one variant being misclassified as homozygotes for the alternative variant.

Most haploblocks were polymorphic (MAF> 5%) in both native (n= 719) and invadad (n=83) regions while only two of them were polymorphic among cultivars (n= 61) (ann1.01 and ann.5.01) (Fig. 3). Based on haploblock allele frequencies, five populations from the invaded range clustered with their putative geographic source while four of them clustered with cultivars (Fig. 3), indicating that at least in some populations, allele frequencies of haploblocks have shifted in the crop direction. We observed no allele frequency shifts in the Texas direction as expected if haploblocks mediated climate adaptation in the invaded range.

**Figure 3.**
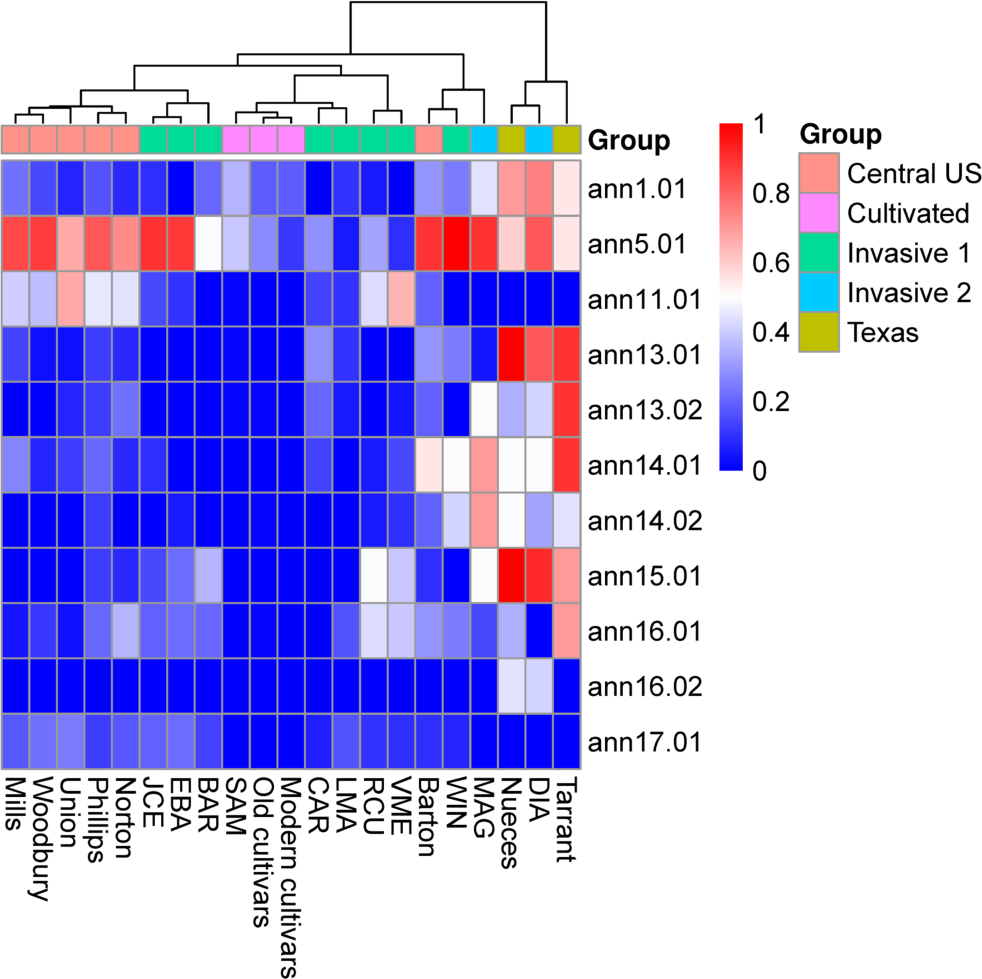
Clustering analysis based on the allele frequency of each of the 11 haploblocks. Clustering was performed without scaling variables and using the Ward D2 method. Heatmap shows the frequency of each haploblock in distinct populations. Populations were grouped as native source (Central US or Texas), invasive (Invasive 1 or Invasive 2), and cultivated groups. The allele frequency of haploblocks for the CROP SAM population was taken from Todesco et al., (2020) and included for comparison with Argentinian cultivars (CROP ARG).

### Genetic divergence post introduction

To identify loci putatively involved in adaptive divergence and invasion success, we performed a genome scan as implemented in *Bayescan*. We performed two comparisons; samples from eight out of 10 invasive populations (all but DIA and MAG, INV1, n= 67) were compared with samples from the six native populations from the central US geographic source (NAT, n= 69) (Fig. 2B) while invasive samples from DIA and MAG (INV2, n= 16) were compared with samples from two Texas populations (TX, n= 20) (Fig. 2C). Outlier loci (q-value< 5%) were found genome-wide in both sets of comparisons (Fig. 4A and 4B). Of the 200 and 55 outliers found in each comparison, respectively, 31 (15.5%) and 11 (20%) were located within haploblocks (significantly more than expected by chance; hypergeometric p = 0.0168 and 0.0096, respectively), and 18 outliers overlapped between comparisons (significantly more than expected by chance; hypergeometric p = 9.59 x 10^-25^). Similarly, of the top 5% of SNPs (based on F_ST_ values) from each comparison, 121 (13.9%) and 113 (14.9%) were located within haploblocks (significantly more than expected by chance hypergeometric p= 0.0006 and 7.54 x 10^-8^, respectively), and 138 overlapped (significantly more than expected by chance; hypergeometric p= 4.51 x 10^-60^). Finally, when haploblocks were treated as single loci, only one haploblock showed strong divergence between NAT and INV1 groups: ann5.01 (F_ST_= 0.19) while three showed moderate to strong divergence between TX and INV2 groups: ann5.01 (F_ST_= 0.17), ann16.01 (F_ST_= 0.33), and ann13.01 (F_ST_= 0.57). The remaining haploblocks, in both sets of comparisons, showed low F_ST_ values between groups in both comparisons (0.0 – 0.09 and 0.0 – 0.07, respectively).

**Figure 4.**
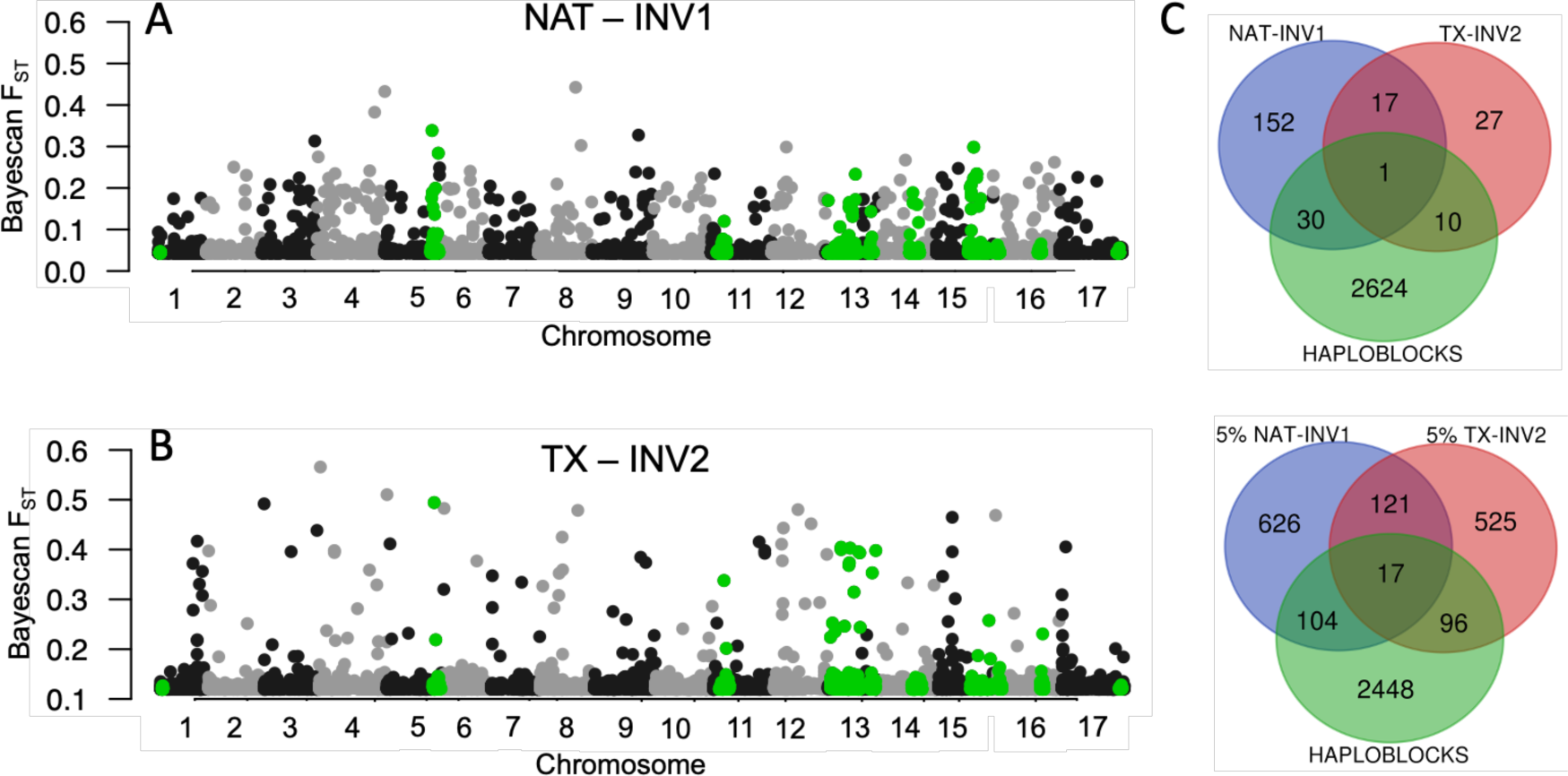
A and B: Bayescan FST between native source and invasive populations, SNPs within haploblock regions are highlighted in green. NAT: 69 individuals from six native populations from the central US (Barton, Phillips, and Norton from Kansas, Mill and Woodbury from Iowa, and Union from South Dakota); TX: 20 individuals from two populations from Texas (Tarrant and Nueces); INV1: 67 individuals from populations sourced from the central region (BAR, CAR, EBA, JCE, LMA, RCU, VME, and WIN); INV2: 16 individuals from populations sourced from Texas (DIA and MAG). C: Venn diagrams showing the overlap between the outliers identified in *Bayescan* (q-value< 5% and top 5% for upper and lower plots, respectively) for NAT-INV1, TX-INV2, and SNPs within haploblocks.

As we detected high levels of admixture between wild *H. annuus* and cultivated sunflower in Argentina, we were interested in the relationship between crop introgression and adaptive divergence. To this end, using the 237 outliers identified in *Bayescan,* we estimated the allele frequency of these outliers in native sources (NAT and TX), invasive (including both INV1 and INV2), and cultivar groups (CROP ARG), and performed an ADMIXTURE analysis to compare the crop ancestry proportions with the one estimated using genome-wide SNPs. If crop alleles contributed disproportionally to genetic divergence, the allele frequency of ‘invasive’ alleles (defined as the most common variant in the invasive group) should be higher in cultivars than native source groups and invasive samples should harbor higher crop ancestry for the outliers than the genome-wide SNPs dataset.

On average, the ‘invasive’ alleles were found at higher frequency in cultivars (median= 0.83) than in the native source groups (median = 0.4 and 0.41 for NAT and TX, respectively) (Fig. 5A). In ADMIXTURE, at K = 2, both genome-wide and outliers datasets separated native and cultivated samples, but invasive and feral samples showed large differences in ancestry proportions between datasets (Fig. 5B and 5C). While individuals from the two native sources showed significantly more crop ancestry for genome-wide than outlier SNPs (median proportion 0.12 and 0.04 for NAT and 0.03 and 0.01 for TX, for genome-wide SNPs and outliers, respectively), the reverse was seen in invasives (median 0.3 and 0.86, respectively) and in the feral population (median 0.66 and 0.92, respectively) (Fig. 5D). All of this indicates that outliers for divergence were substantially enriched by crop alleles.

**Figure 5.**
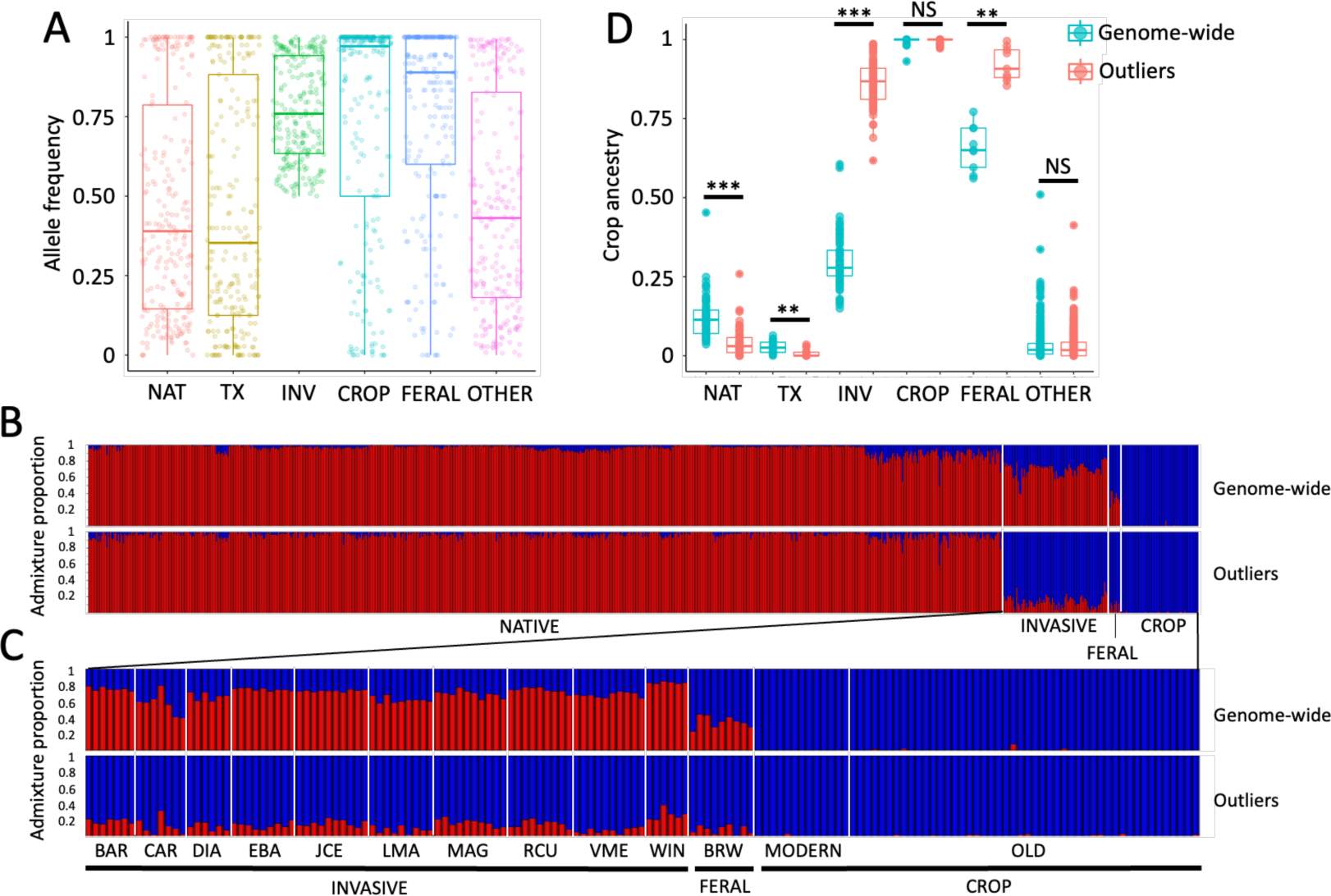
A: allele frequency of ‘invasive’ alleles in distinct groups. For each outlier locus, the invasive allele was defined as the most common variant in the invasive group. B and C: ADMIXTURE analysis at K= 2 for genome-wide SNPs and the 237 outliers identified with *Bayescan*, B) all samples and C) only invasive, feral, and crop samples. D: Median comparisons of crop ancestry proportions between genome-wide and outliers datasets, ***P< 0.0001, **P< 0.01, ^NS^P> 0.05 . NAT: native samples from the central US source; TX: native samples from the Texas source; INV: wild Argentinian *H. annuus* samples; CROP: cultivated samples from Argentina; FERAL: feral samples collected in BRW; OTHER: all native samples not included in either NAT or TX.

## DISCUSSION

Here, we examined the origin and evolution of wild *H. annuus* populations in Argentina. We also investigated whether large haploblocks associated with multiple adaptive traits and climatic variables in the native range showed evidence of selection in the invaded range. We identified two independent geographic sources for Argentinian populations, one in the central US and the other in Texas, and no significant contribution from the related wild annual *Helianthus* species. Multiple genomic regions across the genome showed evidence of selection in Argentina. This was based on allele frequency differences, and was true for both native source-invasive comparions (NAT-INV and TX-INV2). These regions significantly overlapped between comparisons, and showed disproportionally high crop ancestry, indicating that post-introduction admixture with cultivated sunflower strongly influenced the genetic make-up of invasive populations. Finally, two out of 11 haploblocks showed evidence of selection in the invaded range, though for these haploblocks, selection favored the cultivated rather than the texanus haplotype, which is opposite of that predicted by climate adaptation. This implies that the introgression of large structural variants from cultivars may contribute to establishment and spread of introduced species.

### Invasion history

Reconstructing the invasion history, especially identifying the geographic source and the number of independent introductions, is critical for making inferences about post introduction adaptation (Keller and Taylor, 2008; Estoup and Guillemaud, 2010; Sherpa and Després, 2021). Multiple introductions followed by admixture may increase the invasion success of species by alleviating genetic bottlenecks (Dlugosch and Parker, 2008; Dlugosch et al., 2015; Szűcs et al., 2017), creating novel genetic combinations on which selection can act (Ellstrand and Schierenbeck, 2006; van Kleunen et al., 2015), and generating heterozygosity-fitness correlations (Keller et al., 2014; van Kleunen et al., 2015). In this study, we identified two independent geographic sources for invasive populations: the central US, which sourced most of the invasive populations in Argentina, and Texas, source of two invasive populations. This scenario of multiple independent introductions is supported by previous studies based on phenotypic (Cantamutto et al., 2010a) and SSR marker data (Hernández et al., 2019b). The two native sources, while showing low genetic differentiation genome-wide, diverge in ecologically relevant traits as well as at large haploblocks, which are associated with climate adaptation and phenotypic variation (Todesco et al., 2020; Owens et al., 2021). Admixture between these different populations in the invasive range could lead to novel phenotypic and genetic combinations if there is gene flow. However, on average, invasive populations are more differentiated from each other than with native populations (Fig. S1), suggesting little gene flow after dispersal in the invaded range. The lack of gene flow between invasive populations in Argentina has been previously suggested and can be explained by its mosaic distribution, separated in most cases by more than 100 Km as a result of human-mediated dispersal (Cantamutto et al., 2010b). Therefore, while at least two introductions of wild *H. annuus* occurred in Argentina, there is little evidence to support the role of admixture between populations of distinct origin facilitating their establishment and spread.

### Crop introgression

Admixture between cultivated species and their wild relatives is known to have led to the formation of aggressive weeds, capable of invading both natural and agricultural environments (Ellstrand and Schierenbeck, 2000; Hedge et al., 2006; Ellstrand et al., 2010; Bock et al., 2018; Le Corre et al., 2020; Presotto et al., 2023; Vercellino et al., 2023a). Here, we found strong support for admixture and introgression between wild and cultivated sunflowers in the invaded range. These findings align with what has been observed in recent studies on teosinte in Europe (Le Corre et al., 2020), weedy rice in Argentina (Presotto et al., 2023), and wild carrot in the United States (Hernández et al., 2023). The results also suggest that admixture occurred early in the invasion process, before dispersal over the central region of Argentina, because all invasive populations show high levels of crop ancestry, even when several of them have not grown in sympatry with the crop for decades (Poverene et al., 2008; Mondon et al., 2018). Moreover, populations derived from the two independent introduction events showed similarly high levels of admixture, suggesting that admixture occurred at least twice and that crop-wild hybrids are favored in the invaded range. Alternatively, several individuals from the native range (especially from the central US) showed crop ancestry proportions as high as invasive populations; therefore a scenario with introduction of hybrid populations from the native range instead of admixture in the invaded range cannot be ruled out.

Admixture between wild and cultivated individuals may increase invasiveness by distinct non-mutually-exclusive genetic mechanisms including adaptive introgression of large effect loci (e.g., those conferring herbicide resistance; Le Corre et al., 2020; Presotto et al., 2023), transgressive segregation (Campbell et al., 2006; Hedge et al., 2006), and hybrid vigor (Bock et al., 2018; Vercellino et al., 2023b). In our study, most of the genomic divergence between invasives and their native source populations is explained by alleles introduced from cultivars, suggesting that introgression is, at least partially, adaptive. Consistent with this, many traits (e.g., larger disks and seeds, early flowering, and lower seed dormancy) in Argentinian sunflower populations have evolved in the direction of cultivated sunflower (Cantamutto et al., 2010a; Hernández et al., 2019a). It is therefore likely this evolution has been facilitated by the introgression of crop alleles. On the other hand, experimental crosses have shown that early generation crop-wild hybrids in sunflower have low fitness (Mercer et al., 2007; Presotto et al., 2012, 2019), making hybrid vigor an unlikely mechanism. Genetic drift may produce similar patterns to adaptation in genome scans, especially in populations which experienced genetic bottlenecks (Leigh et al., 2021). Genetic bottlenecks are expected to be common during the earliest stages of invasions, and rare alleles that survive the bottleneck can quickly increase in frequency due to genetic drift (Bock et al., 2015; Leigh et al., 2021). Previous studies have shown native and invasive populations harbor similar levels of genetic diversity (Hernández et al., 2019b); however genetic bottlenecks in invasives could have been offset with admixture with cultivated sunflower, and genetic drift cannot explain the bias towards cultivated SNPs. Further work on linking adaptive regions with invasive traits is critical to understand whether crop introgression was causal or incidental to invasion success. In addition, genotyping historical samples and using a temporal approach (Kim et al., 2023) may help elucidate the relative role of genetic drift and adaptation in the evolution of invasiveness in this system.

### No support for climate adaptation driven by large haploblocks

Invasive species are frequently adapted to their local environments yet is unclear whether pre-adaptation to local environments is critical for invasiveness or whether invasive populations are poorly adapted initially and subsequently adapt to local conditions (Colautti and Barrett, 2013; Oduor et al., 2016). In Argentina, invasives established and spread in warmer and less continental environments than one of their native source populations (Cantamutto et al., 2010a; Hernández et al., 2019a). Thus, we hypothesized the haploblocks underlying climate adaptation in the native range (Todesco et al., 2020; Owens et al., 2021) might have facilitated climate adaptation in Argentina as well. While all haploblocks are segregating in the invaded range and several loci within them showed signatures of selection, when haploblocks were treated as single loci, only two of them (ann5.01 and ann13.01) show signatures of selection, though not in the direction predicted by climate adaptation, i.e., increased frequency of the texanus haplotype. Instead, these haploblocks showed evidence of selection in the direction of the cultivar haplotype, consistent with the pattern observed genome wide. One of these haploblocks (ann13.01) is associated with multiple traits that differentiate native and invasive populations (e.g., stem diameter, flowering time, and disk diameter); and could, therefore, have facilitated the evolution of invasive traits. Further studies are needed to understand the phenotypic effects of haploblocks, especially their link with invasive traits.

## ACKNOWLEDGEMENTS

We thank Kaichi Huang for providing support with random forest models, the Biodiversity Research Centre (UBC) for a fellowship to FH, the National Research Council of Argentina (CONICET) for a fellowship to RBV, the Agencia Nacional de Promoción Científica y Tecnológica (ANPCyT) for providing funding (PICT 2019-00722) to FH, and the US Department of Agriculture (USDA) for providing funding to JB for sequencing.

---

**Figure S1.**
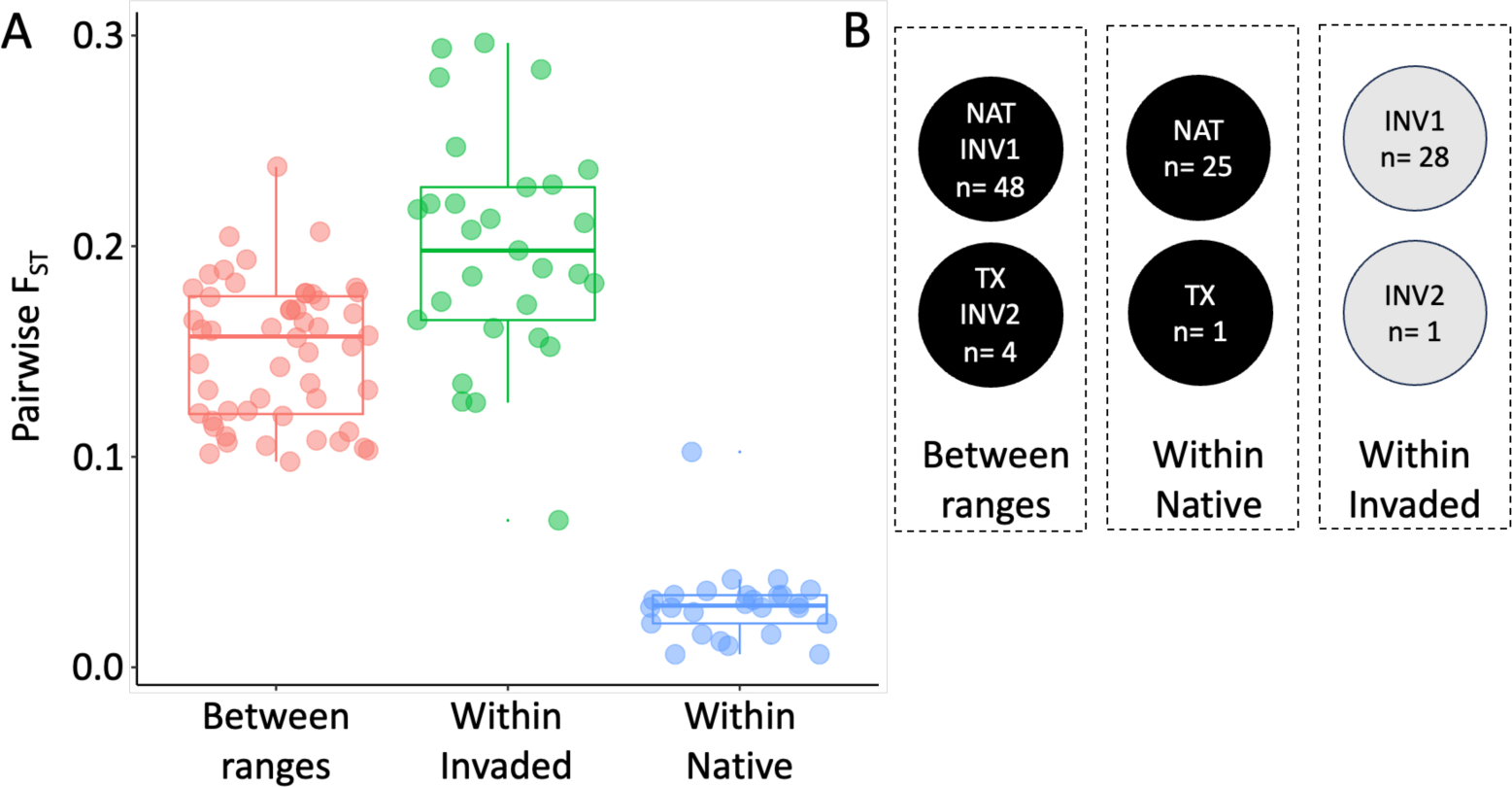
A: Genetic differentiation (pairwise FST) within and between ranges. Populations were assigned to four groups: NAT= 6, TX= 2, INV1= 8, and INV2= 2, and comparisons were performed within groups and between invasive populations and native populations from the putative geographic source (NAT for INV1 and TX for INV2). B: number of comparisons for each category. Median FST of Within invaded was significanty higher than Between ranges (Mann-Whitney Tests: Z= 4.3; P< 0.0001) and Within Native (Z= 6.3; P< 0.0001) categories.

**Figure S2.**
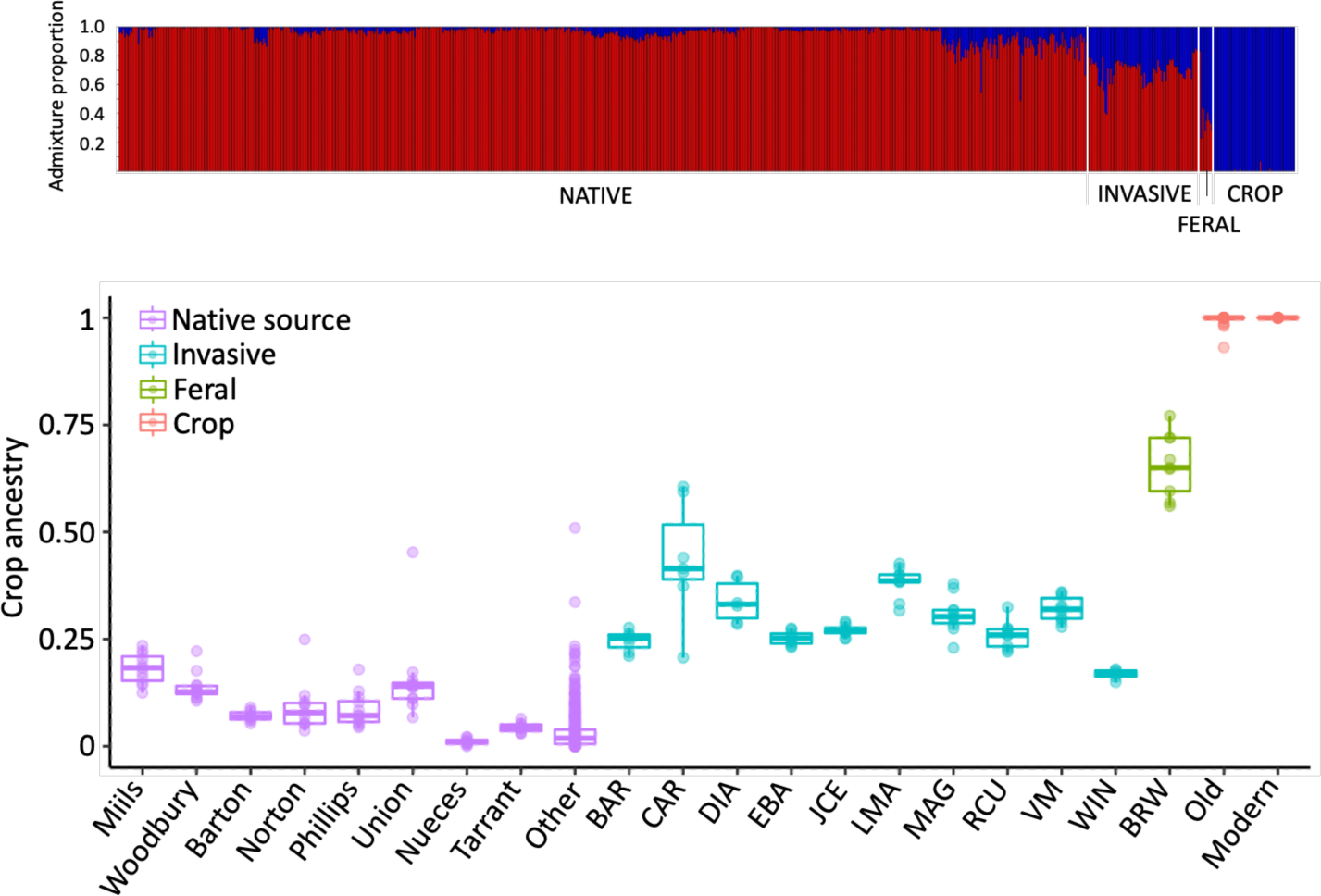
A: ADMIXTURE analysis at K= 2 for genome-wide SNPs. B: crop ancestry proportions for distinct groups; Native source: native populations from the central US (NAT) and Texas (TX) sources; Invasive: invasive populations; Crop: samples from Argentinian old and modern cultivars; Feral: feral samples collected in BRW; Other: all native samples not included in the native source group.

